# Regulation and Characterization of the Cex system of enteric pathogens: a pathway for the lipidation and secretion of noncanoncial bacterial lipoproteins

**DOI:** 10.64898/2026.09.11.750658

**Authors:** Zachary P. Rivas, Kacey M. Talbot, Griffin P. Carter, George P. Munson

**Author notes:** **Correspondence:** George P. Munson.

## Abstract

CexE is an outer membrane lipoprotein of enterotoxigenic *Escherichia coli* (ETEC); one of the most prevalent etiological agents of diarrheal disease. Homologs of CexE are present in enteroaggregative *E. coli*, *Citrobacter rodentium*, and other enteropathogens. We have previously shown that expression of CexE is dependent upon Rns, the master virulence regulator of ETEC and a member of the AraC/XylS superfamily of transcription factors. In this study we report that the expression of *cexD* and *cexPABC*, the genes required for lipidation and delivery of CexE to the outer leaflet of the outer membrane, are also Rns-dependent. Although *cexE* and *cexPABC* are arranged in the same orientation, they are separated by an intergenic region sufficiently large to accommodate a promoter and transcription factor binding sites. Nevertheless, expression of the four gene cluster does not originate within the intergenic region. Rather, *cexPABC* are expressed from the Rns-dependent *cexE* promoter. The remaining gene of the system, *cexD*, is monocistronic and expressed from its own Rns-dependent promoter. Mutagenesis of a predicted Rns binding site upstream of *cexD* abolished Rns binding to a *cexD* promoter fragment in vitro and activation of the promoter in vivo. CexD is a novel bacterial acyltransferase. Unlike canonical bacterial acyltransferases that lipidate amino-terminal cysteines, CexD attaches a lipid to the amino-terminal glycine of CexE. Lipidation must occur in the periplasm because topological mapping revealed that CexD has six transmembrane domains with both its amino-terminus and enzymatic domain in the periplasm. We further show that CexD is required for CexE’s localization to the outer leaflet of the outer membrane and that a *cexD* mutant of *C. rodentium* is attenuated in a murine model. These findings further define the Rns virulence regulon, the enzymology of a noncanonical acyltranferase and its contributions to pathogenicity.

## Introduction

CexE is a lipoprotein that coats the outer leaflet of the outer membrane of enterotoxigenic *Escherichia coli* (ETEC). Homologs are present in enteroaggregative *E. coli* (EAEC), *Yersinia enterocolitica*, *Providencia alcalifaciens,* and the murine pathogen *Citrobacter rodentium* (Hart et al., 2008; Rivas et al., 2020; Sheikh et al., 2002). Using *C. rodentium* as a model enteropathogen we have established that CexE is a virulence factor because the pathogenicity of a *cexE* mutant was significantly attenuated in a neonatal mouse model of infection (Rivas et al., 2020). Furthermore, wildtype *C. rodentium* reached significantly higher loads in the large intestines of adult mice and was shed in greater numbers than a *cexE* mutant (Rivas et al., 2020). The amino-terminus of CexE encodes a signal peptide allowing for its transport to the periplasm via the general secretory Sec-pathway. Removal of the signal peptide by signal peptidase I exposes an invariant amino-terminal glycine (Pilonieta et al., 2007; Rivas et al., 2020; Teufel et al., 2022). Unlike canonical bacterial lipoproteins that are lipidated at amino-terminal cysteines, CexE and its homologs are a newly discovered family of noncanonical lipoproteins in which the lipid is attached to an amino-terminal glycine (Babu et al., 2006; Belmont-Monroy et al., 2020; Icke et al., 2021).

Whereas canonical lipoproteins require Lnt for acylation/lipidation of the exposed cysteine amine, CexE depends upon CexD (AatD) (Gupta et al., 1993; Icke et al., 2021). Two studies have independently shown that CexD is a novel acyltransferase that palmitoylates the amino-terminal glycine of CexE (Dispersin in EAEC) (Belmont-Monroy et al., 2020; Icke et al., 2021). CexD likely obtains palmitoyl from phosphatidylethanolamine; a predominant phospholipid of *E. coli* and other bacteria (Belmont-Monroy et al., 2020; Icke et al., 2021; Raetz, 1978; Rowlett et al., 2017). Four other proteins are required for CexE to reach the outer leaflet of the outer membrane; CexPABC. CexC is a predicted cytosolic ATPase whose activity may be required to extract the nascent lipoprotein from the inner membrane. Consistent with that prediction, we have previously shown that CexE remains in the periplasm of ETEC and *C. rodentium cexC* mutants (Rivas et al., 2020). CexP, a predicted inner membrane protein, may also be involved in the extraction of lipdated CexE from the inner membrane. CexB may shuttle CexE through the periplasm while CexA, a homolog of the outer membrane β−barrel protein TolC, is undoubtedly shuttles CexE across the outer membrane. To increase our understanding of this novel lipoprotein pathway we determined the regulation of the *cex* genes, the subcellular location of CexD’s enzymatic domain, and CexD’s contribution to pathogenicity in competition studies with *C. rodentium* strains in a murine model.

## Results

### Rns-dependent expression of *cexD*

The genetic organization of *cex* and *aat* loci in ETEC, *C. rodentium*, and EAEC have both similarities and differences (Figure 1A). In the prototypical EAEC strain 042 *aatPABCD* are likely transcribed as a polycistronic message while the *cexE* homolog *app* is monocistronic. Based on its location, *cexE* may be monocistronic or bicistronic with the acyltransferase encoding gene *cexD* in *C. rodentium*. In contrast, *cexD* is apparently monocistronic in ETEC. In all three pathogens the expression of the acyltransferase’s client (CexE or Dispersin), is positively regulated by Rns (also known as CfaD or CfaR) or its homologs RegA (C. rodentium) or AggR (EAEC). Rns activates the *cexE* promoter by binding to two experimentally identified sites (Figure 1A) (Pilonieta et al., 2007). To determine if Rns also regulates the expression of *cexD* and other *cex* genes we used a mathematical matrix derived from 31 known Rns binding sites –represented graphically as a DNA logo in Figure 1B– and information theory algorithms to identify and rank potential Rns binding sites (M. D. Bodero et al., 2007; Crooks et al., 2004; Pilonieta et al., 2007; Schneider & Spouge, 1997; Schneider & Stephens, 1990). One highly ranked site was predicted upstream of *cexD* (Figure 1A). Like the promoter proximal sites at *cexEp* and several other Rns activated pilin promoters, the site upstream of *cexD* is in the forward orientation (M. D. Bodero et al., 2008; M. D. R. Bodero & Munson, 2016; G. P. Munson & Scott, 2000; G. Munson & Scott, 1999; Pilonieta et al., 2007).

**Figure 1.**
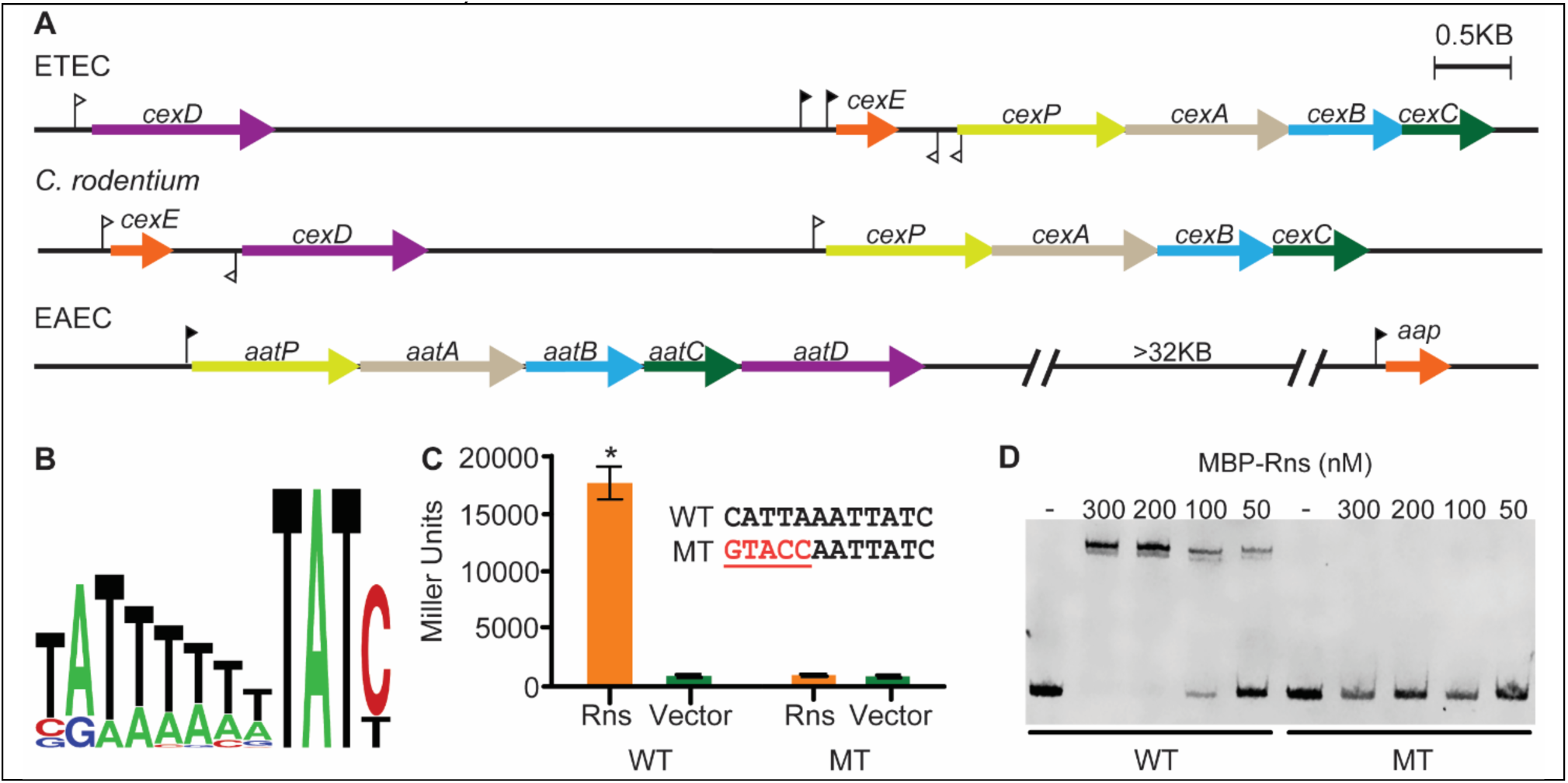
Rns directly activates *cexD* expression. (**A**) Organization of *cex* and *aat* loci in ETEC, *C. rodentium* and EAEC. Homologs are similarly colored and share the same last letter designation with the exception of *aap* which encodes the CexE homolog Dispersin. Unlike in EAEC, the gene encoding the acyltransferase, *cexD,* is not contiguous with *cexPABC* in either ETEC strain H10407 or *C. rodentium* DBS100. Solid and hollow flags denote known and predicted binding sites, respectively, for Rns and its homologs AggR (EAEC) and RegA (*C. rodentium*). The direction of each flag indicates the orientation of the binding site. (**B**) Rns binding site logo derived from 31 known binding sites. (**C**) The *cexD* promoter region was cloned from ETEC H10407 and mutagenized (MT) to construct *cexDp::lacZ* reporter strains. Inset shows the wild-type (WT) and mutagenized Rns binding site with mutations highlighted in red, underlined font. β-galactosidase assays are reported as the mean ± SD. *n* = 3, \**P* < 0.0001 by unpaired Student’s *t*-test. (**D**) EMSA of MBP-Rns with the wild-type and mutagenized *cexD* promoter fragments.

Consistent with the Rns binding site prediction, β-galactosidase assays with Lac reporter strains revealed that Rns activates the *cexD* promoter (Figure 1C). Relative to the vector control, Rns increased β-galactosidase expression 19-fold and the difference was statistically significant. We further confirmed the accuracy of the binding site prediction by oligonucleotide directed mutagenesis. Activation of the *cexD* promoter was abolished by mutagenesis of the predicted binding site (Figure 1C). In vitro, the mutations abolished binding of MBP-Rns to the *cexD* promoter fragment as determined by electrophoretic mobility shift assay (EMSA) (Figure 1D). MBP is used to increase the solubility of Rns for in vitro studies but otherwise has no effect on its activity (G. Munson & Scott, 1999). Collectively these results demonstrate that the expression of *cexD* is Rns-dependent and confirms the existence of an Rns binding site upstream of *cexD* that is required for activation of the *cexD* promoter.

### Rns-dependent expression of *cexPABC*

In addition to the site upstream of *cexD* our algorithm predicted an additional site in the *cexE cexPABC* intergenic region of ETEC as well as second site centered 57 bp inside *cexP* (Figure 1A). In contrast to the site upstream of *cexD,* both of the predicted sites are in the reverse orientation. Although some Rns activated promoters contain binding sites in the reverse orientation, they are invariably accompanied by promoter proximal sites in the forward orientation (M. D. Bodero et al., 2008; M. D. R. Bodero & Munson, 2016; G. P. Munson & Scott, 2000; G. Munson & Scott, 1999; Pilonieta et al., 2007). This observation suggested that two additional sites could be false positives. Nevertheless, we investigated the expression of CexPABC using the terminal gene as a representative of the gene cluster. Due to a lack of antibodies against the native proteins, we tagged CexC with a FLAG epitope and inhibited Rns with decanoic acid. The fatty acid has recently been shown to bind within the amino-terminal domain of Rns and ligand binding abolishes Rns activity (Midgett et al., 2021). As positive and negative controls lysates were also probed for Rns-dependent expression of CexE and DnaK whose expression is Rns-independent. As expected decanoic acid abolished expression of CexE but not DnaK (Figure 2A). Decanoic acid also inhibited the expression of CexC suggesting that expression of CexPABC is also Rns-dependent. To confirm this result we disrupted *rns* with a hygromycin insertion (Δ*rns*). In the Δ*rns* mutant neither CexE nor CexC were produced (Figure 2B). Complementation restored expression of both while the vector control did not. The higher levels of CexE and CexC expression in the complemented strain relative to WT is likely a gene dosage effect produced by the high copy number plasmid used for complementation. These results establish for the first time that expression of CexC –and by inference, CexPAB– is Rns-dependent.

**Figure 2.**
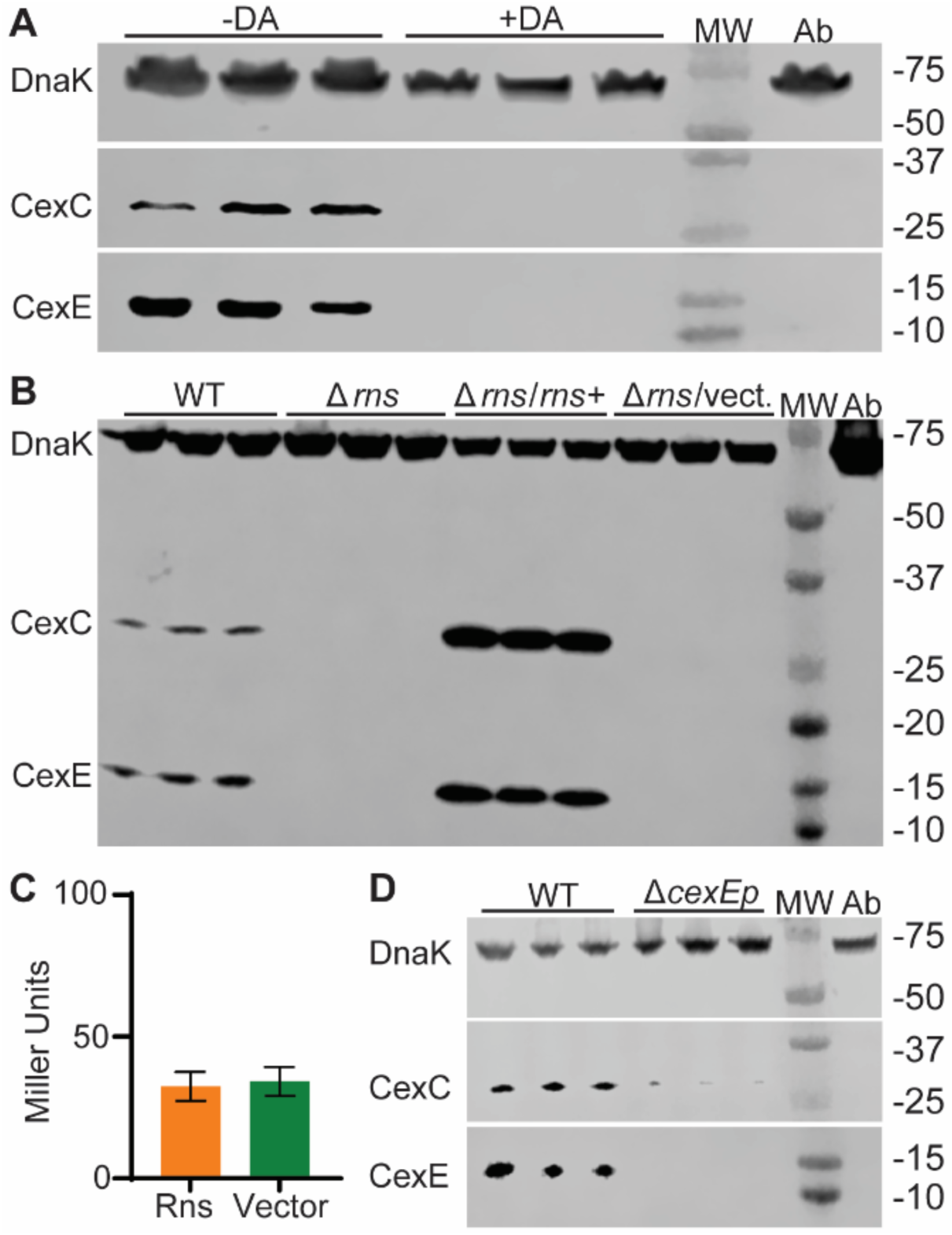
CexPABC expression is Rns-dependent in ETEC. (**A**) Experimental replicates of WT GPM3181 (H10407 *cexC-FLAG*) were cultured with or without 2.5 mM decanoic acid (a fatty acid and inhibitory ligand of Rns) and whole cell lysates were used for Western blot. The same membrane was probed sequentially with antibodies raised against 12 kDa CexE, 26 kDa CexC-FLAG, and 69 kDa DnaK. (**B**) Western blot of GPM3181 and GPM3178 (H10407 *cexC*-*FLAG* Δ*rns*) lysates. The latter was also transformed with a Rns expression plasmid or vector control. Experimental replicates were run in triplicate. The membrane was sequentially probed as described above. (**C**) β-galactosidase assay of a reporter strain with the intergenic region between ETEC *cexE* and *cexPABC* cloned upstream of *lacZ*. The difference between the Rns+ and vector control strain was not significant; *P* = 0.69, Student’s *t* test. *n* = 3, mean ± SD. (**D**) Replicate lysates of GPM3181 and a derivative with the *cexE* promoter deleted. The membrane was sequentially probed as described above. MW, molecular weight ladder; Ab antibody specificity control with lysate of strain GPM1163 (H10407 *cexE::kan*).

We next sought to identify the promoter responsible for Rns-dependent expression of *cexPABC*. To determine if the *cexE cexPABC* intergenic region harbors a Rns-dependent promoter we cloned it into a Lac reporter strain. However there was no difference in β-galactosidase activity between the reporter strain transformed with a Rns expression plasmid or vector control (Figure 2C). Thus, consistent with the atypical orientation of the predicted Rns binding sites, the *cexE cexPABC* intergenic region does not contain a Rns-dependent promoter. We have previously characterized the Rns binding sites upstream of *cexE* and shown that Rns activates its promoter (Pilonieta et al., 2007). To determine if Rns regulates the expression of *cexPABC* through activation of *cexEp* we disrupted the promoter with a hygromycin cassette and probed lysates for CexE and CexC-FLAG expression (Figure 2D). As expected, ablation of *cexEp* abolished CexE expression. Likewise, expression of CexC was abolished (Figure 2D). In aggregate these results indicate that CexE and CexPABC are expressed as a polycistronic message from the Rns-dependent *cexE* promoter.

### Membrane topology of CexD

To determine the subcellular location of CexD’s enzymatic domain we first used the TOPCONS server to predict the topology of the protein; Uniprot ID: E3PPH8 (Tsirigos et al., 2015). The consensus model predicts CexD to be an inner membrane protein with six transmembrane α-helices followed by the enzymatic domain (residues 181-428) oriented in the periplasm (Figure 3A). To assess the validity of the topological model we fused six CexD truncations to PhoA-LacZα to construct dual reporter plasmids (Figure 3A). The enzymatic activity with these reporter plasmids indicates subcellular location as alkaline phosphatase is only active in the periplasm whereas β-galactosidase can only be reconstituted when the LacZα fragment is located in the cytoplasm (Haardt & Bremer, 1996). Fusions to the predicted cytoplasmic loops (R41, S95, R160) produced significantly more β-galactosidase activity relative to the vector control and negligible alkaline phosphatase indicating these fusions are oriented in the cytoplasm (Figure 3B). Conversely fusions to the predicted periplasmic loops (L65, L125) and the enzymatic domain (G198) resulted in significantly higher alkaline phosphatase activity relative to the vector control and negligible β-galactosidase activity (Figure 3B). Collectively these results validate the TOPCONS model and establish that CexD-dependent lipidation of CexE occurs in the periplasm because the enzymatic domain of CexD is periplasmic.

**Figure 3.**
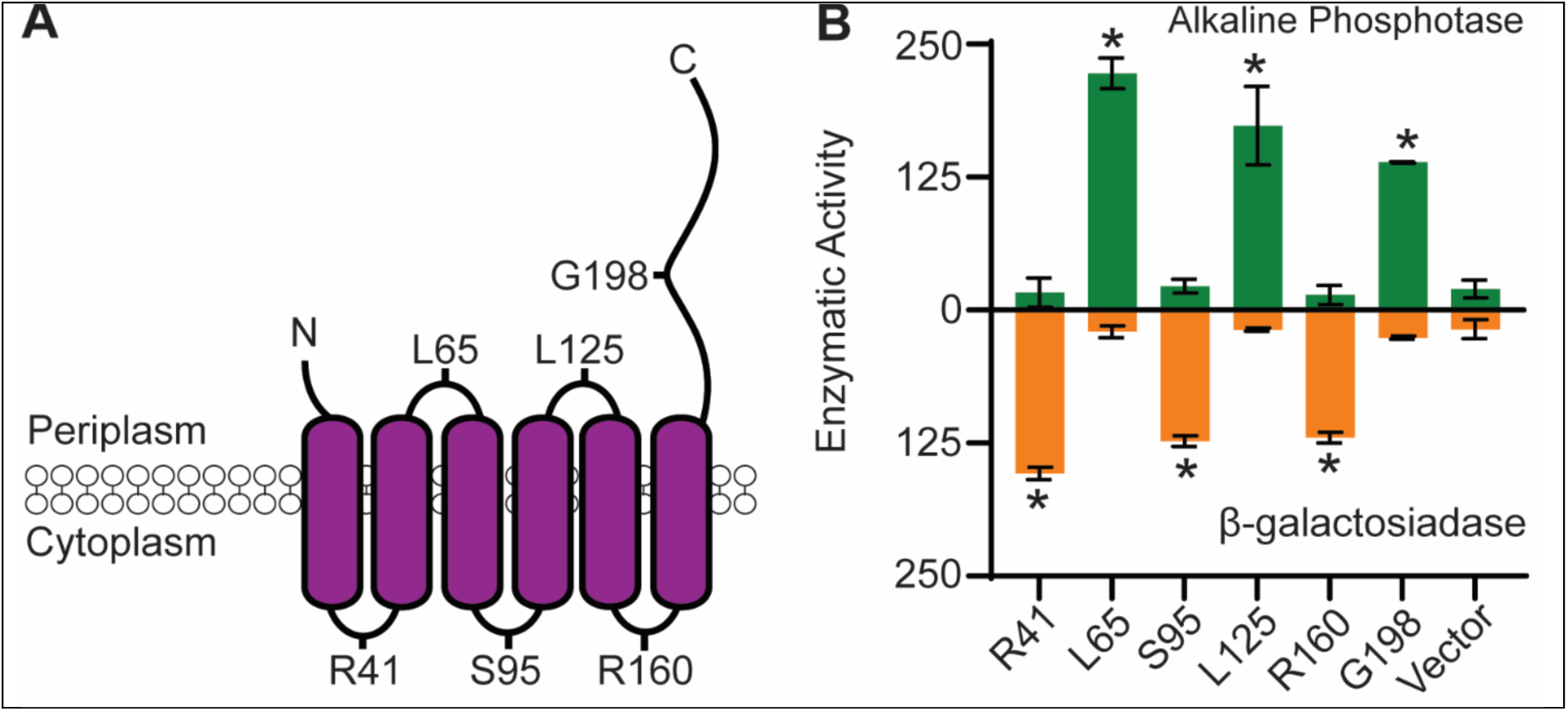
The enzymatic domain of CexD is located in the periplasm. **(A)** Topological model of CexD (Uniprot ID: E3PPH8) from ETEC H10407 as predicted by TOPCONS. CexD is a 47 kDa protein and the first half of the protein contains six predicted transmembrane α-helices. The model further predicts that the carboxy-terminal enzymatic domain is located in the periplasm. Labeled residues denote CexD truncations fused to PhoA-LacZα. (**B**) Strain NEB5α transformed with *cexD::phoA-lacZα* reporter plasmids were used to determine topology based on enzymatic activity. Asterisks denote means that are significantly different (*P* < 0.005 by unpaired Student’s *t*-test) from the vector control, pUC19. *n* = 3, mean ± SD.

### CexD_Cr_ is required for CexE_Cr_ Secretion and increases *in vivo* virulence

We next determined the phenotype of *cexD* mutants in vitro and in vivo. For these experiments we used *C. rodentium* because it is a natural murine pathogen (Barthold et al., 1976; Mundy et al., 2005). To determine if CexD-dependent modification of CexE is required for delivery of CexE to the outer leaflet of the outer membrane we disrupted c*exD_Cr_* and used proteinase K as a membrane impermeable probe (Figure 4A). With WT *C. rodentium* cultured under secretion conditions proteinase digestion of CexE_Cr_ is independent of membrane permeabilization. This indicates that the majority of CexE_Cr_ has been transported across the OM. What little remains in the periplasm is undetectable. In contrast, CexE_Cr_ remains in the periplasm of the *cexD_Cr_* mutant because proteolysis is only apparent after membrane permeabilization (Figure 4A, left image). Complementation of the *cexD_Cr_* mutant restored CexE_Cr_ transport because as with WT, proteolysis was found to be independent of membrane permeabilization (Figure 4A, right image). These results suggest that the Cex secretion system exerts quality control; accepting the modified client for transport across the outer membrane while excluding unmodified CexE. Our results with *C. rodentium* are also consistent with EAEC and ETEC studies (Icke et al., 2021; Nishi et al., 2003).

**Figure 4.**
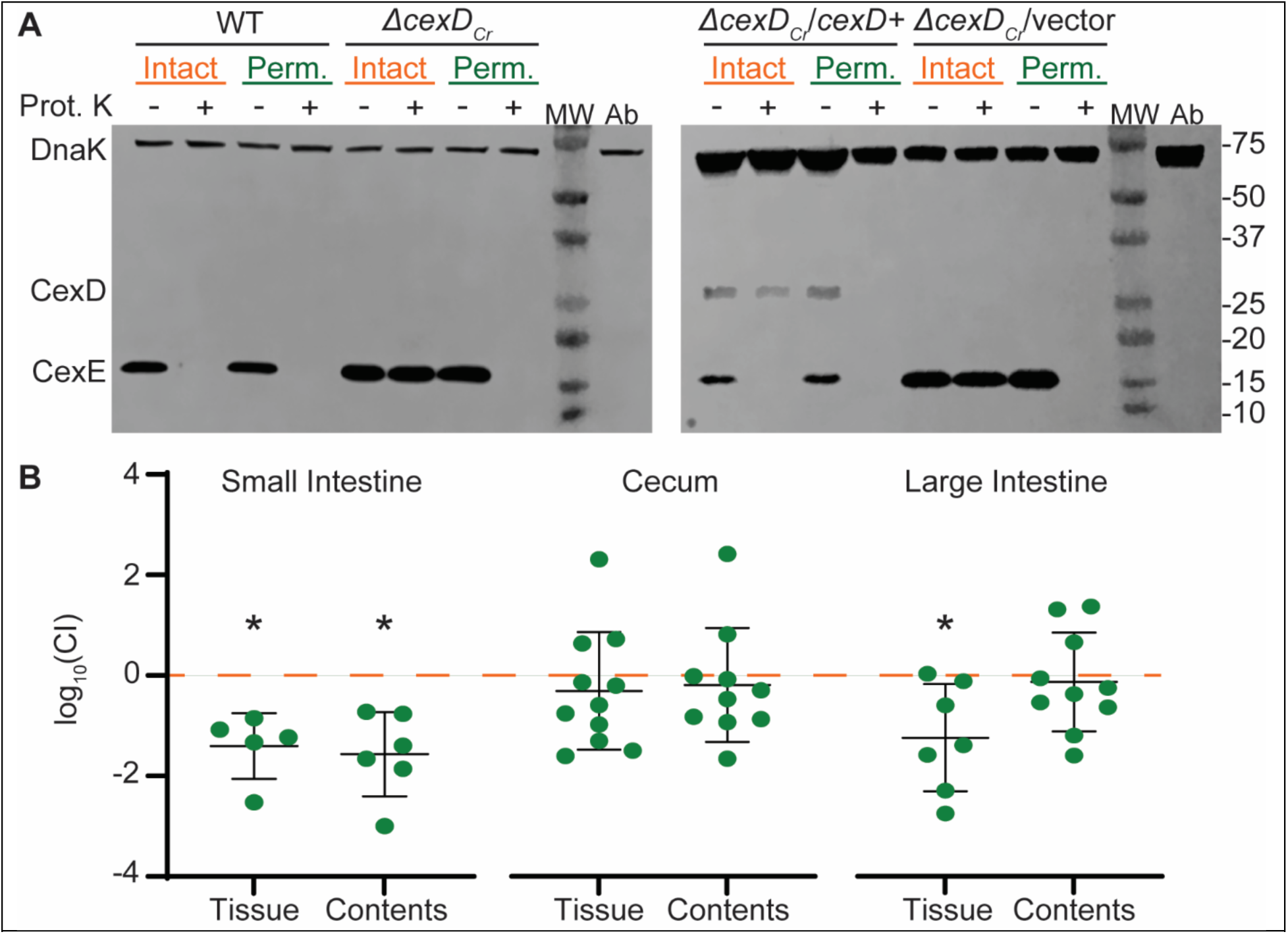
CexD_Cr_ is required for CexE_Cr_ secretion and virulence. (**A**) Left image, membrane permeabilization is not required for enzymatic digestion of CexE_Cr_-FLAG (16kDa) in WT *C. rodentium* strain GPM1830a. In contrast, permeabilization is required for digestion of CexE_Cr_-FLAG in a *cexD* mutant; GPM3174a (DBS100 *cexE_Cr_-FLAG::tet cexD_Cr_::kan)*. Right image, complementation of GPM3174a with a CexD_Cr_-FLAG expression plasmid –but not vector control– restores translocation of CexE_Cr_-FLAG to the outer leaflet of the outer membrane. Although CexD_Cr_-FLAG has a predicted mass of 50 kDa it migrates anomalously at ca. 30kDa. Membranes were probed for the FLAG epitope then DnaK. Representative blots shown, *n* = 3. Prot. K, proteinase K; MW, molecular weight ladder; Ab, antibody specificity controls. Antibody specificity control lanes were loaded with whole cells lysates of *C. rodentium* strain DBS100 which lacks the FLAG epitope. (**B**) Competitive index (CI) of intestinal loads of C57/Bl6 mice orogastrically inoculated with either WT *C. rodentium* or *cexD_Cr_* mutant GPM3180 and GPM1391-A1, respectively. Intestinal tissue and contents were harvested 16 days post-inoculation. CFUs normalized to sample mass and log_10_ transformed. Dotted line represents a log_10_ value of 0 which would indicate equal recovery of WT and *cexD_Cr_* mutant strains. Mean ± SD are shown, asterisk indicates the mean is significantly different from 0, *n* = 5-11 mice per group, \**P* < 0.05 by one sample t-test.

To determine the impact of CexD in vivo we co-inoculated C57Bl/6 mice via oral-gastric gavage with our WT *C. rodentium* strain –which carries an *aadA* cassette in the intergenic region downstream of *cexD_Cr_*– and the *cexD_Cr_::kan* mutant used in our proteinase K digestion assays. At sixteen days post inoculation the WT strain was found to have significantly outcompeted the mutant in the small intestine tissue and contents as well as the large intestine tissue (Figure 4B). Although statistical significance was not achieved with the contents of the large intestine nor cecum samples, in all cases a trend of greater WT fitness than mutant was observed. In aggregate these results establish that CexD-dependent palmitoylation of CexE contributes to the pathogenicity of *C. rodentium* and confirm our previous in vivo studies with a *cexE* mutant (Rivas et al., 2020).

## Discussion

We have previously shown that the expression of the outer membrane lipoprotein CexE is Rns-dependent (Pilonieta et al., 2007). Here we have extended that work to elucidate the expression of the five genes required for CexE’s lipidation and transport to the outer membrane’s outer leaflet. In H10407 and other ETEC strains the gene encoding the acyltransferase, *cexD*, is apparently monocistronic and we have found that Rns activates expression of *cexD* through occupancy of a site upstream of the gene. Although *cexE* was also expected to be a monocistronic gene we have found that it is expressed in a polycistronic message with *cexPABC*. This was surprising because *cexE* and *cexPABC* are separated by a 272 bp intergenic region. However, Lac reporters demonstrated that the intergenic region lacks a Rns-dependent promoter even though inhibition of Rns abolished expression of CexC. Subsequent mutagenesis of the *cexE* promoter abolished expression of CexE and CexC. Thus, *cexEPABC* are apparently expressed as a polycistronic message that originates at the Rns-dependent *cexE* promoter.

Although microarray data has shown that RegA is required for expression of *cexE* and *cexPABC* in *C. rodentium*, the genes are unlikely to be polycistronic given their spacing and arrangement (Figure 1A) (Hart et al., 2008). Rns and RegA have 46% identity between their DNA binding domains, can complement one another, and recognize similar DNA binding sites (Hart et al., 2008; G. P. Munson & Scott, 2000). Our analysis has identified a putative Rns/RegA binding site upstream of *C. rodentium cexPABC* that may be involved in RegA-dependent expression of the four genes. Microarray data from *C. rodentium* strain ICC168 -which has a similar arrangement of *cex* genes to DBS100-has shown that *cexD* is also positively by RegA (Hart et al., 2008). Although a Rns/RegA binding site is predicted in the *cexE cexD* intergenic region, its reverse orientation suggest the site is not functional. Indeed we found this to be the case for the *cexE cexPABC* intergenic region of ETEC (Figures 1A and 2). Our results with ETEC raise the possibility that in *C. rodentium cexD* is expressed as a bicistronic message from a RegA-dependent promoter upstream of *cexE*.

CexD is responsible for the palmitoylation of CexE and it has been hypothesized that CexD obtains palmitoyl from phosphatidylethanolamine; a phospholipid that comprises 70-80% of the *E. coli* inner membrane (Belmont-Monroy et al., 2020; Raetz, 1978). Not surprisingly, CexD is a predicted inner membrane protein and we have mapped its six transmembrane α-helices which comprise the amino-terminal half of the protein. CexD has an N– and C–out topology thus placing its enzymatic carboxy-terminal domain in the periplasm. Although TOPCONS predicts that EAEC AatD (Uniprot ID: D3H597) and *C. rodentium* CexD (Uniprot: D2TJZ6) have just five transmembrane α-helices, in both cases the enzymatic domains are predicted to be periplasmic with the acyltransferases having N–in, C–out topology. Because CexE remains in the periplasm of *cexD* mutants (Figure 4A), lipidation of CexE is a prerequisite for its secretion across the periplasm and outer membrane. Therefore, there is a quality control checkpoint, likely residing in one or more of the other Cex proteins, responsible for ensuring that CexE has been properly modified prior its delivery to the outer leaflet.

This work and our previous study have established that CexE must be exposed on the outer leaflet of the outer membrane to function as a virulence factor. In vivo we have found that a *cexD* mutant of *C. rodentium* is attenuated and outcompeted by a WT strain (Figure 4B). Likewise, we found that a *cexC* mutant was attenuated in a murine enteric infection model (Rivas et al., 2020). CexC is a predicted cytosolic ATPase that may be required for the extraction of lipidated CexE from the inner membrane. Consistent with this prediction CexE remains in the periplasm of *cexC* mutants (Rivas et al., 2020). Although not yet investigated, it is likely that CexE is an inner membrane lipoprotein in *cexC* mutants because CexD is capable of lipidating CexE in the absence of the other Cex proteins (Belmont-Monroy et al., 2020; Icke et al., 2021). In addition to questions regarding the Cex secretion machinery, it is also not yet clear how CexE contributes to pathogenicity. In EAEC Dispersin has been proposed to disrupt bacterial aggregation through electrostatic repulsion of pili (Velarde et al., 2007). However this hypothesis is an extrapolation from the overexpression of Dispersin from the strong T5 promoter and therefore may be an artifact. It has also been shown that CexE coats outer membrane vesicles (Roy et al., 2011). This raises the possibility that CexE may contribute to pathogenesis through a mechanism independent of pili. While the precise mechanism of CexE remains under investigation this work clarifies CexD’s contribution to CexE’s acylation, secretion, and pathogenesis.

## Acknowledgements

The authors thank Zoe Thomas for technical assistance.

Research reported in this publication was supported by the National Institute of Allergy and Infectious Diseases of the National Institutes of Health under award numbers R21AI128164 and R01AI168157. The content is solely the responsibility of the authors and does not necessarily represent the official views of the National Institutes of Health.

## Material and Methods

### Bacterial Growth Conditions

Bacteria were cultured aerobically at 37°C in in Iscoves Modified Dulbecco’s Medium (IMDM; Thermo Fisher Scientific) with or without 2.5mM decanoic acid in 0.4% DMSO. For hypoxic growth conditions bacteria were cultured in sealed CyroELITE^TM^ cyrogenic vials (Wheaton) at 37°C in IMDM. Growth medium was supplemented with 50 µg / ml kanamycin, 150 µg / ml ampicillin, or 5 µg / ml tetracycline as appropriate.

### Bacterial Strains and Plasmids

Strains, plasmids, and primers sequences are listed in Tables 1-3, respectively. HiFi DNA assembly was used to construct plasmid pCexD_Cr_-FLAG which expresses CexD_Cr_-FLAG (∼50kDa) from the pBAD promoter. The CexD_Cr_ vector backbone was amplified from pBADTags2 with primers 1415/1416. *CexD_Cr_* was amplified from *C. rodentium* strain DBS100 genomic DNA with primers 1679/1680. HiFi was also used to sequentially construct cloning vectors pTags2-hyg and pTags2-hyg-rgnB. The pTags2-hyg vector backbone was amplified from pTags2 with primers 1533/1534. The hygromycin gene was amplified from pVitro2-hygro-MCS with primers 2081/2082 and PCR products were assembled with NEB HiFi. The plasmid pTags2-hyg-rgnB vector backbone was amplified from pTags2-hyg with primers 2288/2289. The *rgnB* gene was amplified from pHKLac1z with primers 2290/2291. λRED mediated recombineering was used for the construction of the following strains as previously described (Datsenko and Wanner, 2000; Datta et al., 2006). The kanamycin cassette for epitope tagging *cexC* was amplified from pSUB11 with primer pair 1145/1146. Electroporation of the cassette into H10407 / pSIM6 resulted in strain GPM3181. Hygromycin resistance cassettes targeting the *cexEp* in ETEC H10407 were amplified from pTags2-hyg and pTags2-hyg-rgnB with primer pairs 2125/2126 and 2292/2293, respectively. Electroporation of the cassette into GPM1820a / pSIM6 resulted the recombinants GPM3178 and GPM3179a, respectively. The cassettes for disruption or insertion downstream of *cexD_Cr_* were amplified from pAH120 and pAH144 with primer pairs 1442/1444 and 1482/1483, respectively. Electroporation of the cassettes into DBS100 / pSIM6 resulted in strain GPM1391-A1 and GPM3180, respectively. The cassette for epitope tagging *cexE_Cr_* was amplified from pSUB11-tet with primers pair 1178/1179. Electroporation of the cassette into GPM1391-A1 / pSIM5 resulted in strain GPM3174a. Recombinants were cured of λRED expression plasmids by passage at 42°C. Insertions were verified by PCR with primers flanking the insertion sites.

**Table 1:** Strains.

| **Strain** | **Characteristics** | **Notes** |
| --- | --- | --- |
| NEB5α | *E. coli* K-12 *fhuA2* $\Delta$(*argF-lacZ*)U169<br>*phoA glnV44* Φ80 $\Delta$(*lacZ*)M15 *gyrA96*<br>*recA1 relA1 endA1 thi-1 hsdR17* | (Anton & Raleigh, 2016) |
| H10407 | ETEC O78:H11 CFA/I+ ST+ LT+ CexE+ | (Dolores G. Evans et al., 1975) |
| DBS100 | *Citrobacter rodentium* (ATCC 51459) | (Schauer et al., 1995) |
| MC4100 | *E. coli* K-12 F- *araD139* $\Delta$(*argF-lac*)U169<br>*rpsL150 (StrR) relA1 flhD5301 deoC1*<br>*ptsF25 rbsR* | (Casadaban, 1976) |
| KS1000 | *F' lacIq lac+ pro+ / ara del(lac-pro)*<br>*del(tsp)=del(prc)::KanR*<br>*eda51::Tn10(TetR) gyrA rpoB thi-1*<br>*argI(am)* | (M. D. Boderio et al., 2007) |
| GPM1391-A1 | DBS100 *cexD<sub>Cr</sub>::kan* | This study |
| GPM1830a | DBS100 *cexE<sub>Cr</sub>-FLAG::kan* | (Rivas et al., 2020) |
| GPM3159 | MC4100 *attB<sub>HK022</sub>::pCexPLac1z* | This study |
| GPM3174a | DBS100 *cexE<sub>Cr</sub>-FLAG::tet cexD<sub>Cr</sub>::kan* | This study |
| GPM3178 | H10407 *cexC-FLAG::kan rns::hyg* | This study |
| GPM3179a | H10407 *cexC-FLAG::kan cexEp::hyg* | This study |
| GPM3180 | DBS100 *cexD<sub>Cr</sub>::kan, cexE<sub>Cr</sub> $\Omega$*<br>(114bp::*aadA*) intergenic *aadA* insertion<br>downstream of *cexD<sub>Cr</sub>::kan* | This study |
| GPM3181 | H10407 *cexC-FLAG::kan* | This study |
| GPM3182 | MC4100 *attB<sub>HK022</sub>::pCexDLac1z* | This study |
| GPM3183 | MC4100 *attB<sub>HK022</sub>::pCexDLac2z* | This study |

**Table 2:** Plasmids.

| **Name** | **Description** | **Marker** | **Notes** |
| --- | --- | --- | --- |
| pGPMRns-Myc | Rns-myc cloned into pTags2 expressed from *lacp* | *bla* | (Midgett et al., 2021) |
| pNEB193 | Cloning vector | *bla* | New England Biolabs |
| pUC19 | Cloning vector | *bla* | Addgene |
| pSIM5 | $\lambda$ RED expression plasmid | *cat* | (Datta et al., 2006) |
| pSIM6 | $\lambda$ RED expression plasmid | *bla* | (Datta et al., 2006) |
| pAH120 | PCR template for *kan* cassette | *kan* | (Haldimann & Wanner, 2001) |
| pAH144 | PCR template for *aadA* cassette | *aadA* | (Haldimann & Wanner, 2001) |
| pSUB11-tet | PCR template for tet cassette | *tet*,<br>*bla* | *NIH Figshare* |
| pTags2 | Cloning vector | *bla* | NIH FigShare |
| pVitro2-MCS | Cloning vector | *hyg* | Invivogen |
| pTags2-hyg | Cloning vector | *bla*,<br>*hyg* | NIH Figshare |
| pTags2-hyg-rgnB | Cloning vector | *bla*,<br>*hyg* | This study |
| pRARE2 | Provides seven rare *E. coli* tRNAs | *cat* | Novagen |
| pMBPRns | MBP-Rns expressed from IPTG-inducible pTac promoter | *bla* | (M. D. Boder et al., 2007) |
| pBADTags2 | Cloning vector | *cat* | *NIH Figshare* |
| pCexD<sub>Cr</sub>-FLAG | CexD<sub>Cr</sub>-FLAG cloned into pBADTags2 expressed from *pBAD* promoter | *cat* | This study |
| pCexDp193 | *cexDp* (-125 to +125 relative to ORF) cloned into pNEB193. | *bla* | This study |
| pHKLac1z | Integration plasmid | *aadA* | (Haldimann & Wanner, 2001) |
| pCexDpLac1z | *cexDp*(-125 to +125 relative to ORF) cloned into pHKLac1z | *aadA* | This study |
| pCexDpLac2z | *cexDp*(-125 to +125 relative to ORF) with five point mutations: -103C to G, -102A to T, -101T to A, -100T to C, and -99A to C cloned into pHKLac1z. | *aadA* | This study |
| pCexPpLac1z | *cexDp*(-291 to +166 relative to ORF) cloned into pHKLac1z | *aadA* | This study |
| pPHO1 | Cloning vector | *bla* | (Silva-Herzog et al., 2008) |
| pPhoALac | Cloning vector | *bla* | This study |
| pCexD<sub>R41</sub> | *cexD* (+1 to +123 relative to ORF) cloned into pPhoALac expressed from *lacp*. | *bla* | This study |
| pCexD<sub>L65</sub> | *cexD* (+1 to +195 relative to ORF) cloned into pPhoALac expressed from *lacp*. | *bla* | This study |
| pCexD<sub>S95</sub> | *cexD* (+1 to +285 relative to ORF) cloned into pPhoALac expressed from *lacp*. | *bla* | This study |
| pCexD<sub>L125</sub> | *cexD* (+1 to +375 relative to ORF) cloned into pPhoALac expressed from *lacp*. | *bla* | This study |
| pCexD<sub>R160</sub> | *cexD* (+1 to +480 relative to ORF) cloned into pPhoALac expressed from *lacp*. | *bla* | This study |
| pCexD<sub>G198</sub> | *cexD* (+1 to +594 relative to ORF) cloned into pPhoALac expressed from *lacp*. | *bla* | This study |

**Table 3:**
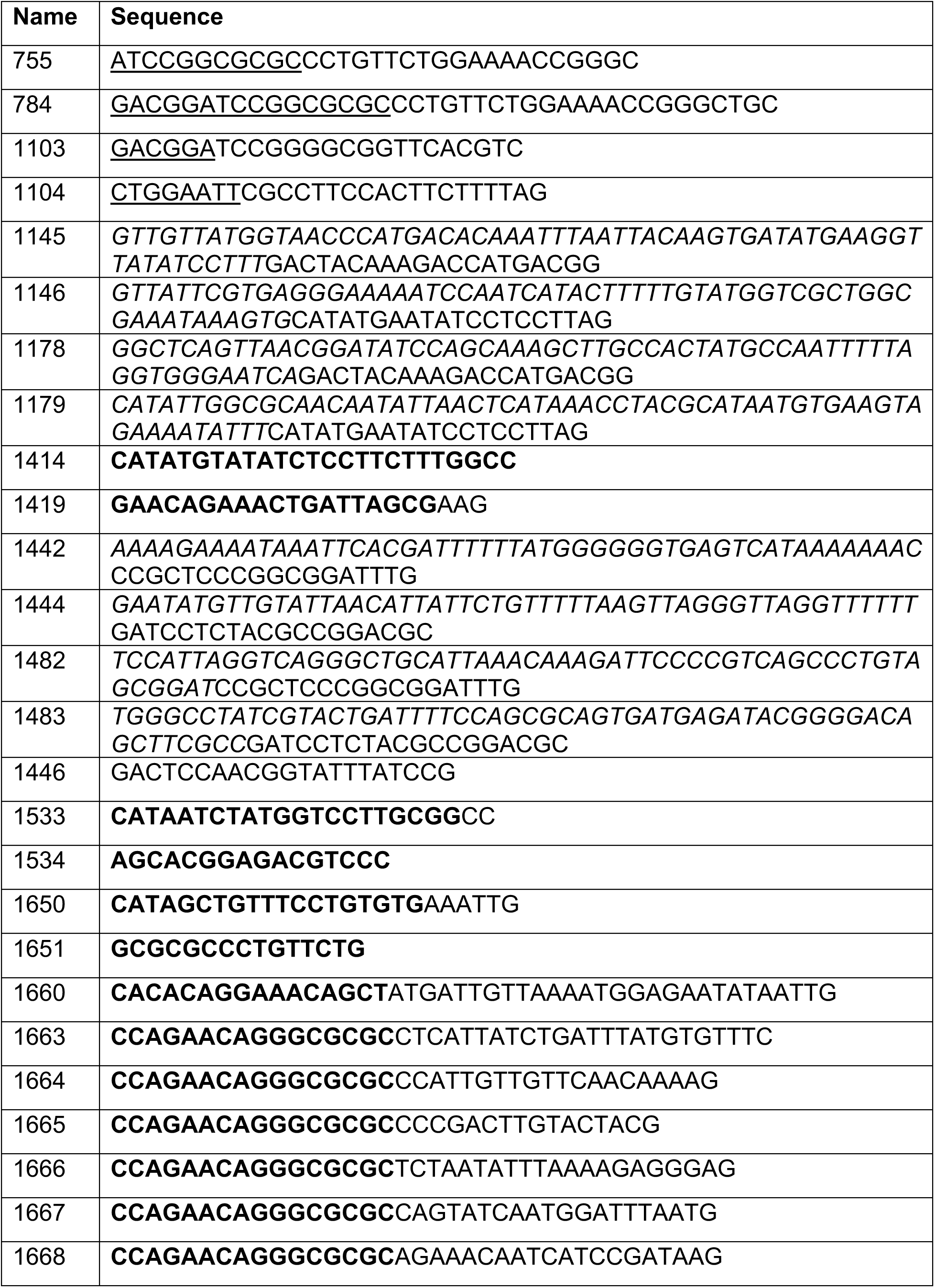
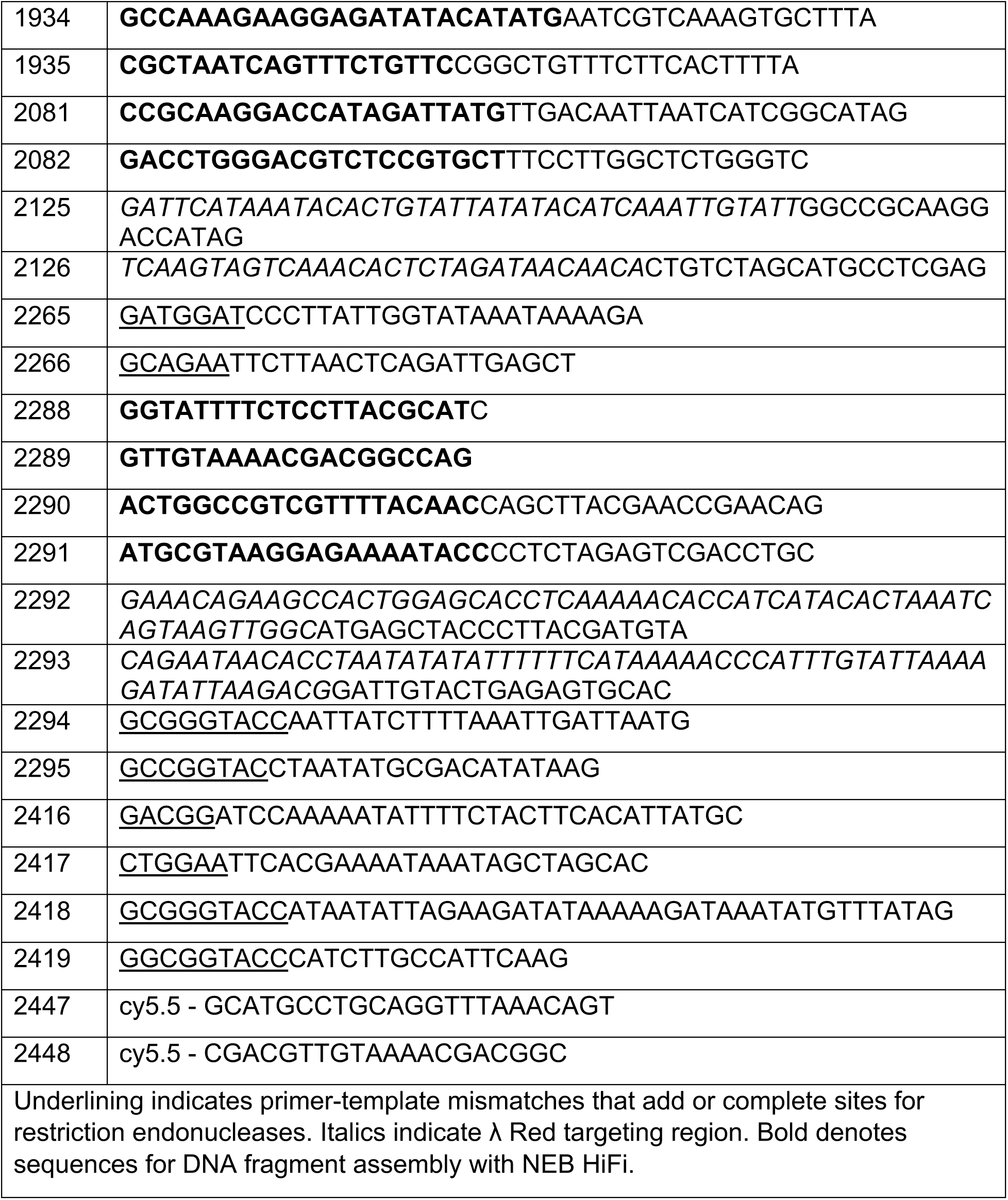
Oligonucleotides.

### Construction of β-galactosidase reporter strains

The plasmid pHKLac1z is a promoterless *lac* reporter plasmid carrying the *pir*-dependent R6Kγ origin of replication, *aadA* conferring resistance to spectinomycin and streptomycin, and *attP*_HK022_. Transcriptional terminators are located upstream and downstream of *lacZ* to prevent readthrough from flanking sequences. The CexP promoter was amplified from ETEC strain H10407 with primers 2265 and 2266. The CexD promoter was amplified from H10407 with primers 1103 and 1104. The PCR products were digested with BamHI-HF and EcoRI-HF (New England Biolabs) and ligated into the same sites of the pHKLac1z vector backbone to construct pCexPLac1z [*cexPp*(−291 to +166) relative to ORF)::*lacZ*] and pCexDLac1z [*cexDp*(−125 to +125 relative to ORF)::*lacZ*]. Additionally the *cexDp* PCR product was also cloned into the BamHI and EcoRI sites of pNEB193 (NEB) to construct pCexD193. Plasmid pCexD193 was subjected to oligonucleotide-directed mutagenesis by primers 2294 and 2295 to introduce point mutations within the Rns binding site upstream of *cexDp.* The PCR products were then digested KpnI-HF (NEB) and ligated using T4 DNA ligase (NEB). The point mutations were confirmed by Sanger sequencing and the mutagenized promoter fragments were cloned into the BamHI-EcoRI fragment resulting in pCexDLac2z. Plasmid pCexDLac2z carries the same *cexDp* with the exception of five point mutations −103C to G, −102A to T, −101T to A, −100T to C, −99A to C. Each reporter plasmids was integrated into the chromosome of MC4100 [F^−^ *araD139* Δ(*argF-lac*)*U169 rpsL150 relA1 flhD5301 deoC1 ptsF25 rbsR*] as previously described to produce strains GPM3159 (*attB*_HK022_::pCexPLac1z), GPM3182 (*attB*_HK022_::pCexDLac1z), GPM3183 (*attB*_HK022_::pCexDLac2z) and single integration was confirmed by colony PCR (Haldimann & Wanner, 2001).

### β-Galactosidase assays

Reporter strains GPM3159, GPM3182, and GPM3183 were transformed with pGPMRns-myc (Rns+) or a vector plasmid pTags2 and were grown aerobically at 37°C in LB with 150µg/ml ampicillin. Cells were harvested during log growth phase, lysed, and assayed for β-galactosidase activity as previously described (Miller, J. H.1972. Experiments in molecular genetics. Cold Spring Harbor Laboratory, Cold Spring Harbor, NY.).

### Expression and purification of MBP-Rns

Maltose binding protein (MBP) fused to the amino terminus of Rns was expressed and purified from a protocol modified from (M. D. Bodero et al., 2007). KS1000/pRare2/pMBPRns1 was grown aerobically at 37°C in LB broth, containing 30 μg/ml chloramphenicol, and 150 μg/ml ampicillin. After cells reached mid-log phase, the culture was transferred to a 25°C shaking water bath, and expression of MBP-Rns was induced by the addition of 500 μM IPTG. After overnight induction, cells were harvested at 4°C and concentrated >100-fold in ice-cold lysis buffer A (20 mM Tris-Cl (pH 7.6), 200 mM NaCl, 1 mM EDTA). Cells were lysed by two passages through a French press. Insoluble material was removed by centrifugation of the lysate at 40,000 × *g* for 40 min at 4°C. The supernatant was loaded on a 1 ml MBPTrap HP column (Cytiva). The MBP-Rns fusion protein was eluted from the column with buffer B (buffer A with 10 mM maltose). Fractions containing MBP-Rns were loaded on a HiPrep 26/10 Desalting column (Cytiva) and eluted in 10 mM Tris-Cl (pH 7.4), 280 mM NaCl, 1 mM EDTA, 10 mM β-mercaptoethanol.

### Electromobility shift Assay

The 5’ cyanine 5.5 labeled DNA fragments were generated by PCR using primers 2447/2448 with pCexDp193 as the template. 2.5 nM of the labeled *cexD* promoter fragments were incubated with 50-300 mM purified MBP-Rns at room temperature for 60 min in binding buffer (10 mM Tris-Cl (pH 7.4), 50 mM KCl, 1mM DTT, 1 ng/µl poly(dI-dC), and 100 µg/ml BSA. Glycerol was added to a final concentration of 6.5% (v/v) prior to separation on 5% native polyacrylamide gels. Gels were analyzed with the Odyssey FC Imaging System (LI-COR Biosciences).

### Construction of *cexD::phoAlacZα* reporter plasmids

The dual *phoA-lacZα* gene fusion reporter system was used to determine the membrane topology of CexD in ETEC H10407 (Alexeyev & Winkler, 1999). Plasmid pPHO1 encoding the *E. coli phoA* minus its N-terminal secretion signal was amplified with primers 755 and 784 (Silva-Herzog et al., 2008). The 1.3kB PCR product was digested with *Asc*I and *Kpn*I and ligated into the same sites in pNEB193 generating plasmid pPhoALac. The plasmid pPhoALac has an IPTG-inducible promoter upstream of the native portion of the *E. coli* alkaline phosphatase and the α peptide fragment of *E. coli β*-galactosidase. The pPhoALac vector backbone was amplified by primers 1650 and 1651. The CexD insert DNA fragments were generated by PCR with a shared forward primer 1660, and a unique reverse primer 1668, 1667, 1666, 1665, 1664, 1663 to amplify DNA encoding the CexD residues + 1 to + 41, + 1 to + 65, +1 to + 95, +1 to +125, +1 to +160, +1 to +198, respectively. HiFi DNA assembly was used to construct pCexD_R41_, pCexD_L65_, pCexD_S95_, pCexD_L125_, pCexD_R160_, pCexD_G198_. Confirmed correct insertions by sanger sequencing.

### Alkaline phosphatase / β-galactosidase assays

The dual PhoA/LacZα reporter plasmids, pCexD_R41_, pCexD_L65_, pCexD_S95_, pCexD_L125_, pCexD_R160_, pCexD_G198_, were electroporated into *E. coli* NEB5α to determine alkaline phosphatase and β-galactosidase activity. NEB5α / pUC19 was used as a negative control. Alkaline phosphatase activity was measured using a protocol modified from (Haardt & Bremer, 1996; Karimova & Ladant, 2017). Cells were grown in LB containing the appropriate antibiotics at 37°C with aeration to an OD_600_ of 0.2-0.3 and the cultured was induced by adding 1mM IPTG and grown at the same conditions until an OD_600_ of 0.7-0.8. 200µl of the culture was pelleted and washed in an equal volume solution of 10mM Tris-HCl (pH 8.0) and 10mM MgSO_4_. Cells were pelleted and resuspended in an equal volume solution of 1M Tris-HCl (pH 8.0), 0.1mM ZnCl_2_, and 1mM iodoacetamide. 50µl of 0.1% SDS and 50µl of chloroform were added to permeabilize cells. Reactions were initiated by the addition of 1mM *p*-nitrophenyl phosphate and terminated by the addition of 2M sodium hydroxide. Enzymatic activity was calculated using the method previously describe (Miller, J. H. Experiments in Molecular Genetics (Cold Spring Harbor Laboratory Press, 1972).

### Proteinase K Digests

Bacteria were collected from 600 μl of log phase cultures by centrifugation at 21,000 g for 1 min. and resuspended in 75 μl of either PBS or PBS, 10 mM EDTA, 1% (vol/vol) Triton X-100. Suspensions were equilibrated at 37°C for 1 h prior to addition of proteinase K to a final concentration of 200 μg/ml. Samples were incubated for 1 h at 37°C with continuous gentle rocking. Enzyme activity was terminated by addition of PMSF followed by 25 μl of 4x SDS-PAGE loading buffer with β-mercaptoethanol.

### Immunoblots

Whole-cell lysates were subject to SDS-PAGE, transferred to PVDF membranes, and blocked in TBS-Blotto (25 mM TrisCl pH 7.6, 150 mM NaCl, 5% (wt/vol) powdered nonfat milk. Antibodies against CexE were produced and used as previously described (Pilonieta et al., 2007; Rivas et al., 2020). Anti-FLAG (A00187) was purchased from GenScript and used at a dilution of 1:5,000. Anti-DnaK (ab69617) was purchased from AbCam and used at a dilution of 1:10,000. Conjugated goat anti-rabbit (sc-2030) and goat anti-mouse (115-036-062) antibodies were purchased from Santa Cruz Biotechnology and Jackson ImmunoResearch Laboratories, respectively. Secondary antibodies were used at a dilutions of 1:10,000. Chemiluminescence was detected with an Odyssey FC Imaging System (LI-COR Biosciences).

### In vivo studies

C57Bl/6 mice were purchased from Jackson Laboratories and housed under barrier conditions in the Division of Veterinary Recourses at the University of Miami Miller School of Medicine. 8 week old mice were orogastrically inoculated with a 200µl of a 1:1 ratio of *C. rodentium* strains (GPM1391-A1) and (GPM3180) in PBS using a 22-gauge, round-tipped feeding needle. Administered CFUs were determined by serial dilution and plating of inocula. At day 16 post inoculation mice were euthanized and their organs were homogenized in cold PBS using an OMNI International Tissue Homogenizer (Kennesaw, GA, United States) for 2 min at medium speed. Homogenates were diluted and plated on MacConkey agar plates with kanamycin or spectramycin/spectomycin. The competitive index was calculated as the quotient of the *cexD_Cr_* mutant to the WT *C. rodentium*. One sample t-test was used to determine if the mean was significantly different from a log_10_ value of 0.

